# Mid-infrared chemical imaging of living cells enabled by plasmonic metasurfaces

**DOI:** 10.1101/2024.09.17.613596

**Authors:** Steven H. Huang, Po-Ting Shen, Aditya Mahalanabish, Giovanni Sartorello, Jenny Li, Xuefeng Liu, Gennady Shvets

**Affiliations:** Cornell University, School of Applied and Engineering Physics, Ithaca, NY, 14853, USA; The Ohio State University, Department of Pathology, Comprehensive Cancer Center, College of Medicine, Columbus, OH, 43210, USA

**Keywords:** infrared microscopy, vibrational microscopy, metasurfaces, plasmonics, surface-enhanced infrared absorption (SEIRA)

## Abstract

Mid-Infrared (MIR) chemical imaging provides rich chemical information of biological samples in a label-free and non-destructive manner. Yet, its adoption to live-cell analysis is limited by the strong attenuation of MIR light in water, often necessitating cell culture geometries that are incompatible with the prolonged viability of cells. Here, we introduce a new approach to MIR microscopy, where cells are imaged through their localized near-field interaction with a plasmonic metasurface. Chemical contrast of distinct molecular groups provided sub-cellular resolution images of the proteins, lipids, and nucleic acids in the cells that were collected using an inverted MIR microscope. Time-lapse imaging of living cells demonstrated that their behaviors, including motility, viability, and substrate adhesion, can be monitored over extended periods of time using low-power MIR light. The presented approach provides a method for the non-perturbative MIR imaging of living cells, which is well-suited for integration with modern high-throughput screening technologies for the label-free, high-content chemical imaging of living cells.

## 1 Introduction

Mid-infrared (MIR) micro-spectroscopy has become a powerful technique for vibrational imaging of biological samples due to its ability to analyze samples non-destructively and without labels, leveraging the unique molecular vibrational fingerprints for chemical specificity.^1,2^ Traditionally, IR microscopy has been performed with Fourier-Transform Infrared (FTIR) spectrometers coupled with focal plane array (FPA).^3,4^ Recently, the advancement of high-brilliance, widely tunable quantum cascade lasers (QCLs) has also popularized discrete-frequency (DF) IR microscopy.^5,6^ Applied to biological samples, IR microscopy has been used to image dried and thinly sliced tissue samples for the label-free histological assessments of diseases.^3,4,7,8^ Alternatively, employing small, IR-active vibrational probes such as azides, deuterium, and ^13^C, mid-infrared (MIR) imaging has demonstrated remarkable potential in the metabolic profiling of cells and tissues.^9,10^

Despite the potential of MIR microscopy in biomedical analysis, MIR microscopy is poorly suited for studying live cells in culture because the MIR light is strongly attenuated in water. For MIR transmission measurements, cells typically need to be placed in a thin cuvette (∼10 μm), but such tight confinement could interfere with the long-term cellular behavior and proliferation.^11^ Attenuated total reflection – Fourier transform infrared (ATR-FTIR) micro-spectroscopy can image cells in reflection mode, but it requires the cells to be cultured on ATR-crystals, complicating its scaling to high-throughput microwell-based geometries.^4^ While MIR photothermal imaging greatly improves the resolution of MIR imaging by using a visible probe beam to detect the heat generated by the MIR pump, the small thermo-optic coefficient of water (on the order of 10^-4^/K) leads to weak photothermal signal from cells.^12–16^ Raman scattering is another complementary vibrational imaging technique, but spontaneous Raman imaging requires high optical intensity due to the small Raman cross section, and coherent Raman imaging such as stimulated Raman scattering (SRS) microscopy suffer from limited spectral coverage and the need for expensive instruments.^17–19^

Here, we introduce a novel approach to the label-free MIR imaging of cells in culture – Metasurface-enabled Inverted Reflected-light Infrared Absorption Microscopy (MIRIAM) – that utilizes the surface-enhanced infrared absorption (SEIRA) effect^20,21^ achieved when analyte molecules are placed in the hotspots of plasmonic nanoantennas^22,23^ arranged into MIR-resonant metasurfaces. When the vibrational fingerprints of the analyte and metasurface resonances spectrally overlap, the local modulation of the nanoantennas scattering cross-sections by the analyte can be detected in the reflected far field. Previously, such metasurfaces have been used for the detection and monitoring of self-assembled monolayers and biomolecules such as protein monolayers and lipid bilayers through IR spectroscopy.^24–28^ Our group has also integrated MIR metasurfaces with cell culture chambers and used them for monitoring dynamic changes in living cells, including cholesterol depletion, cellular response to therapeutics, and cytoskeletal reorganization in response to activation of signaling pathways.^29–31^ Recently, the application of MIR metasurface in the imaging of dried tissue thin section samples has been demonstrated to have increased sensitivity to ultrathin tissue regions.^32^ However, the application of MIR metasurfaces to the imaging of living cells – in particular, with vibrational contrast and sub-cellular resolution capable of visualizing distinct organelles – has not been shown so far.

In our approach, cells are cultured on a MIR metasurface attached to the bottom of a microwell cell culture chamber,^31^ and MIR images of cells on the metasurface are acquired in the reflection mode using an inverted QCL-based scanning confocal microscope: see Fig. 1 for a schematic and Fig. S1 for details. For hydrated fixed cells, hyperspectral data cubes were sequentially obtained as two-dimensional (2D) images at DFs, enabling the imaging of the cellular organelles (nuclei, cytoplasm, and lipid droplets) distinguished by the MIR vibrations of their predominant molecules – nucleic acids, proteins, and lipids, respectively. Additionally, the MIR “movies” of living cells in culture medium were acquired as time-lapse images at one or several wavenumbers, enabling the observation of dynamic cellular processes such as cell spreading and locomotion.

**Fig. 1.**
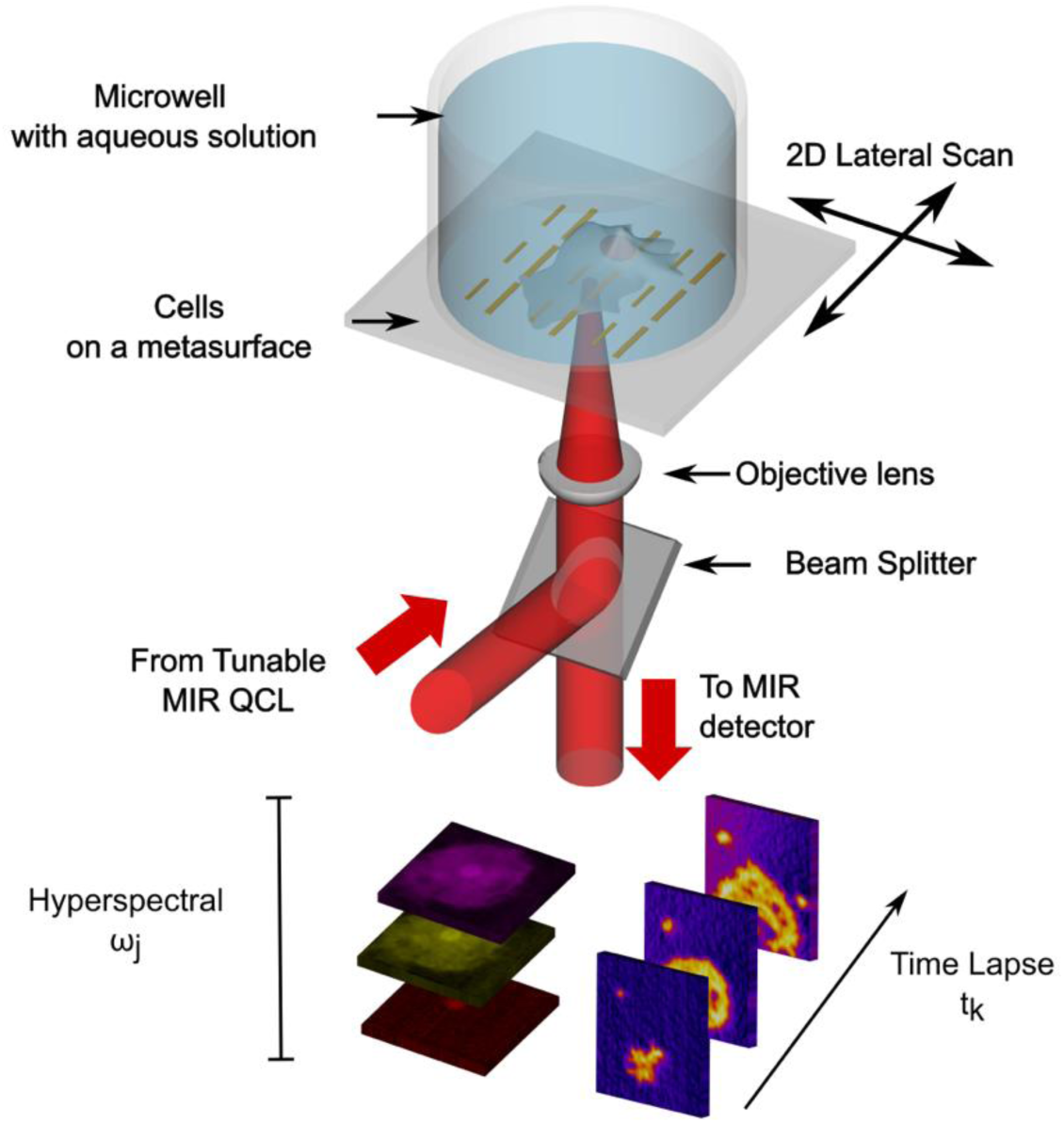
Schematic of the Metasurface-enabled Inverted Reflected-light Infrared Absorption Microscopy (MIRIAM). Cells are cultured in a metasurface-bottomed cell culture chamber and imaged from the bottom using a tunable MIR QCL. As no optics is required above the sample, the cell culture can be conveniently manipulated directly on the platform from above. Hyperspectral and time-lapse images of the cells are formed using the intrinsic chemical contrast of different biomolecules in the cells.

## 2 Materials and Methods

### 2.1 MIR microscope

An inverted MIR confocal laser scanning microscope (modified based on an upright microscope design reported previously^7^ was developed to accommodate the use of metasurface integrated with microwells for live cell imaging. Detailed schematics of the optical and electronics setups are presented in Fig. S1. The emission from a tunable QCL (MIRcat-1400, Daylight Solutions) was passed through an OD2 attenuator and a CaF_2_ beam-splitter, then focused onto the metasurface from below. A Black Diamond-2 infrared laser collimation lens with NA = 0.71 (390093, LightPath Technologies) was used as the focusing objective. The IR beam reflected by the metasurface was collected by the same objective, reflected by the CaF_2_ beam-splitter, and passed through a 100 μm confocal aperture to eliminate any stray light reflected from outside the focal plane. Finally, the IR light was detected using a liquid-nitrogen-cooled MCT detector (MCT-13-1.0, InfraRed Associates).

The signal from the MCT detector was amplified through a pre-amplifier (MCT-1000, InfraRed Associates) and a lock-in amplifier (SR830, Stanford Research Systems). The output from the lock-in amplifier was read by a data acquisition card (PCIe-6361, National Instruments). The QCL pulse width was set to 100 ns for the protein/lipid band (1,500 - 1,800 cm^-1^) and 980 ns for the phosphate band (1,050 - 1,150 cm^-1^). The repetition rate was set to 100 kHz. The time constant of the lock-in was set to 300 µs. The QCL has a varying power across the tuning range; for the QCL module used for the protein/lipid band, the peak average power was measured to be 14 µW at λ = 6.7 µm just before entering the objective. For the QCL module used for the phosphate band, the peak average power was 26 µW at λ = 9.1 µm. The laser polarization was fixed to vertical polarization from the QCL, corresponding to the long axis of the nanoantennas (See Fig. S3).

To form images, the metasurface sample was loaded on a dual-axis motorized microscope stage (HLD117, Prior Scientific), and the sample was scanned in a snake scan pattern. The maximum scan speed along the x-direction was 2 mm/s and the average scan speed was approximately 1 mm/s for a typical image. We used 1 µm pixel size to characterize the PSF of our system using PMMA nano-blocks and 2 µm pixel size for every other measurement. An offset to the x position was added to the backward scans of the snake scan to correct for imaging artifacts related to a spatial delay introduced by the lock-in amplifier.

### 2.2 Metasurface *fabrication*

The plasmonic nanoantennas were patterned by electron beam lithography (JBX9500FS, JEOL) on a 240 nm-PMMA-coated 12.5 mm × 12.5 mm × 0.5 mm IR-transparent CaF_2_ substrate (Crystran). An anti-charging agent (DisCharge H_2_O, DisChem) was spun before exposure. After developing the pattern with 1-to-3 methyl isobutyl ketone (MIBK): isopropanol, 5 nm of chromium and 70 nm of gold were deposited, and the nanoantennas were formed by lift-off in acetone. The metallic nanoantennas formed an array covering a 500 µm × 500 µm area (the metasurface). See Fig. S3 for the detailed design of the nanoantennas. A gold stripe of 500 µm ×30 µm was placed on each side of the metasurface for background measurement.

To image the metasurface in water or culture medium, the metasurface was attached to the bottom of a cell culture chamber superstructure (CS16-CultureWell Removable Chambered Coverglass, Grace Bio-Labs).

### 2.3 PMMA nano blocks

PMMA nano-blocks were patterned onto the single-resonant metasurface to investigate the imaging principle in our reported technique. Using aligned exposure with electron beam lithography (JEOL JBX9500FS), arrays of 2.2 µm × 600 nm × 240 nm PMMA blocks were fabricated, with a periodicity of 27 µm in both the horizontal and vertical directions. These PMMA nano-blocks were aligned with gold nanoantennas in three different configurations. In the first group, each PMMA nano-block was aligned to be positioned directly on top of a gold nanoantenna. In the second group, each PMMA nano-block was aligned to be positioned in the middle between two gold nanoantennas. In the third group, PMMA nano-blocks were placed on bare CaF_2_ substrates without any gold nanoantennas nearby.

### 2.4 Cell Culture

#### 2.4.1 3T3-L1 Pre-adipocytes (fibroblasts)

Mouse embryonic fibroblast cell 3T3-L1 (acquired from the American Type Culture Collection) were cultured in Dulbecco’s Modified Eagle Medium containing 4.5 g/L glucose and GlutaMAX supplement (DMEM, Gibco), further supplemented by 10% fetal bovine serum (FBS, Gibco) and 1% penicillin/streptomycin (P/S, Gibco) in standard incubator condition (37°C, 5% CO_2_). To transfer cells to metasurface, cells were trypsinized using 0.25% Trypsin-EDTA (Gibco) and seeded on metasurface coated with 10 μg/mL fibronectin (Sigma-Aldrich). Cells were incubated overnight to allow for cell adhesion and spreading on metasurface.

For hydrated fixed cell measurement, cells on metasurface were fixed with 4% formaldehyde (Invitrogen) for 15 minutes and washed three times with phosphate-buffered saline (PBS, Gibco). The fixed cells were permeabilized by 0.5% (v/v, in PBS) Triton X-100 (Sigma-Aldrich) for 15 minutes. Then, cells were stained with 300 nM DAPI stain (Invitrogen) for 5 minutes, washed three times with PBS, followed by 6.6 µM Alexa 488 phalloidin stain (Invitrogen) for an hour, then washed three times with PBS.

For live cell measurement, 3T3-L1 cells were trypsinized from culture flask and re-suspended in Leibovitz’s L-15 Medium (L15, Gibco), supplemented with 10% FBS and 1% Antibiotic-Antimycotic (Gibco). Cells were seeded on metasurface coated with 0.2% gelatin (Sigma-Aldrich) just before the start of the measurement. Live cells were imaged at 20 °C and ambient atmospheric condition.

#### 2.4.2 3T3-L1 Adipocytes

Pre-adipocyte 3T3-L1 cells (passage < 10) were grown in DMEM, supplemented with 10% FBS and 1% penicillin/streptomycin until confluency. After another 48h incubation of the confluent cells, the culture medium was exchanged to differentiation medium consisting of DMEM supplemented with 10% FBS, 1% P/S, 1 μM dexamethasone (Sigma-Aldrich), 500 μM 3-isobutyl-1-methylxanthine (IBMX, Sigma-Aldrich), 2 μM rosiglitazone (Sigma-Aldrich), and 2 μg/mL bovine insulin (Sigma-Aldrich). After three days incubation in differentiation medium, the culture medium was exchanged to adipocyte maintenance medium consisting of DMEM supplemented with 10% FBS, 1% P/S, and 2 μg/mL bovine insulin. Cells were incubated in adipocyte maintenance medium for another 7 days. At this point, cells were trypsinized from culture flask and re-seeded on metasurface coated with 0.2% gelatin and incubated in adipocyte maintenance medium for another 5 days. Upon sufficient maturation of lipid droplets, cells were fixed with 4% formaldehyde for 15 minutes and washed in PBS three times.

### 2.5 Image Acquisition and Processing

For imaging PMMA nano-blocks, the absorbance image of PMMA nano-blocks on nanoantenna (Fig. 2c) is presented with no post-processing, whereas the absorbance image of PMMA nano-blocks between nanoantennas (Fig. 2f) was Gaussian-smoothed to enhance the visibility of the PMMA pattern. To obtain the absorbance spectrum shown in Fig. 2(g), the mean of six pixels centered on the PMMA image and six background pixels were used.

**Fig. 2:**
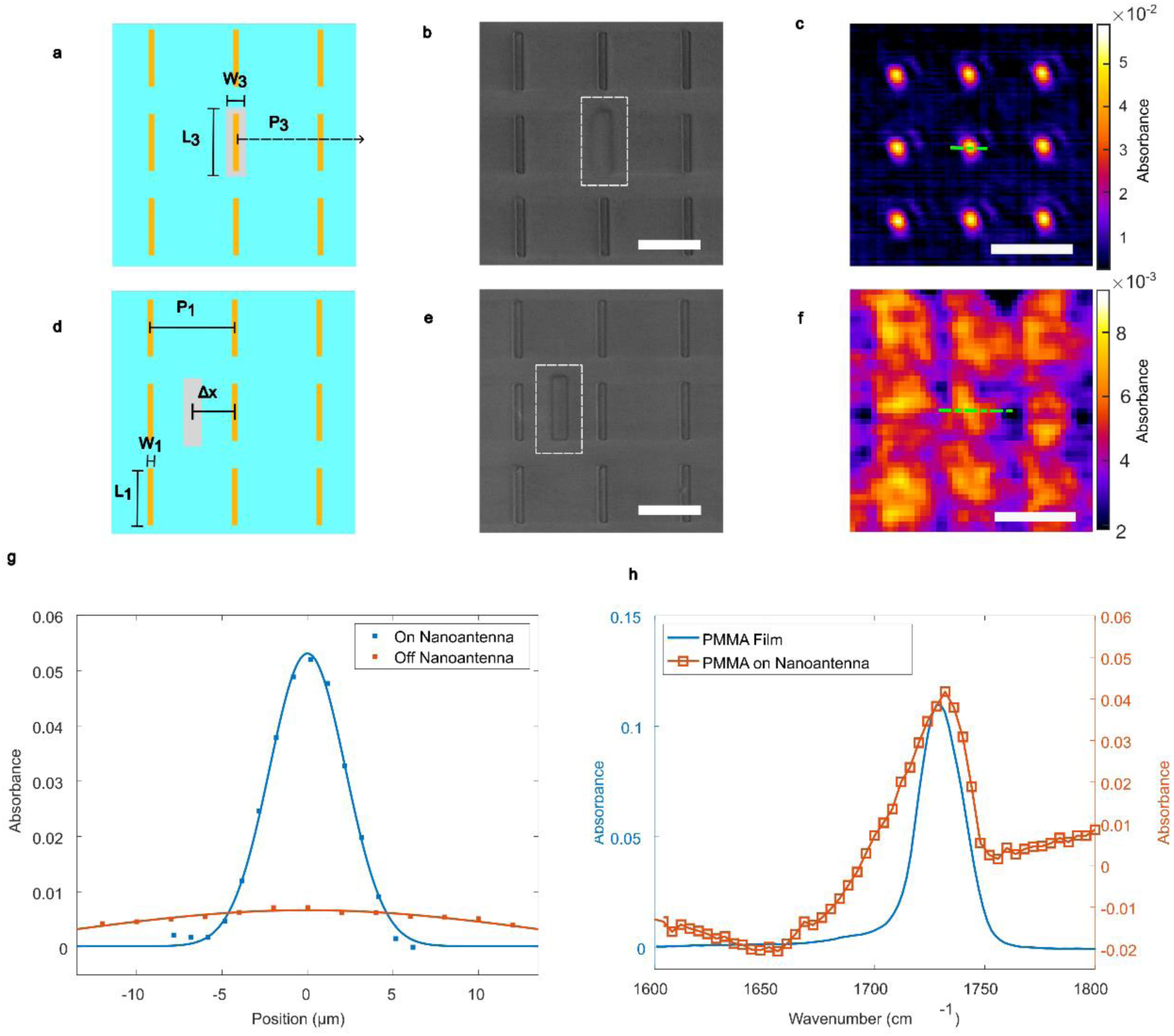
Imaging an array of PMMA blocks on metasurface immersed in H_2_O using the MIRIAM platform. (a-f) PMMA blocks and their MIR images: on (a-c) and between (d-f) the nanoantennas. Schematic drawing (a,d) and the SEM images of PMMA blocks inside the white dotted boxes (b,e). Scale bar: 2 µm. PMMA blocks are arranged as a super-array with ×10 periodicity of the nanoantennas. (c) and (f) shows the MIRIAM absorbance images of the PMMA blocks at 1,730 cm^-1^. Scale bar: 30 µm. (g) Intensity profile of the MIR images along the green dotted lines in (c,f): blue (c) and red (f). On nanoantenna: FWHM = 5.23 µm. Off-nanoantenna: FWHM = 33.67 µm. (h) MIRIAM-based absorbance spectra of the PMMA blocks on nanoantennas in H_2_O (red line) compared to FTIR absorbance of a thin PMMA film in air. Geometrical parameters: L1 = 1.8 µm, W1= 200 nm, P1 = 2.7 µm, L3 = 2.2 µm, W3 = 600 nm, P3 = 27 µm, ***Δx*** = ***P***_**1**_/**2**.

For imaging fixed cells, DF images were acquired in the 1,500-1,800 cm^-1^ range, at 5 cm^-1^ spacing, for the amide I/II vibrations of proteins and ester carbonyl (C=O) vibration of lipids. For imaging the symmetric PO_2_^−^ phosphate stretching, images were acquired in the 1,050-1,130 cm^-1^range, at 5 cm^-1^ spacing. The raw signal was first normalized by the signal at a large patch of gold beside the metasurface (akin to background measurement in conventional FTIR spectroscopy) to obtain a reflectance image. Next, a 2D low pass filter was applied to the image to remove artifacts related to nanoantenna periodicity. This is necessary when employing the bi-resonant nanoantenna design because a grid-like pattern emerges when imaged at high wavenumbers. This effect occurs because the periodicity of the nanoantennas becomes comparable to the size of the focus spot of the MIR beam. The use of 2D low pass filter is optional for the single-resonant nanoantenna design. A flat field correction is applied to correct for illumiation non-uniformity. Savitzky–Golay filter is then applied to smooth the spectra. Then, the reflectance images *R*(*x*, *y*) are converted to absorbance images 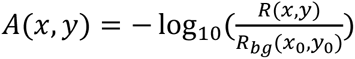, where *R*_b g_(*x*_0_, *y*_0_) is the background reflectance at a position within the metasurface that has no cell. The hyperspectral image cube is then processed with minimum noise fraction (MNF) for noise reduction.

Finally, baseline subtraction is applied to obtain images with vibrational contrast. We have chosen to use a linear baseline, using the images at 1,500 cm^-1^ and 1,800 cm^-1^ to compute the baseline for amide I/II vibrations and ester carbonyl vibration. Images at 1,070 cm^-1^ and 1,100 cm^-1^ are used to compute the baseline for the PO_2_^-^ phosphate vibration. This baseline value is subtracted from the DF images corresponding to each vibration (amide I: 1,655 cm^-1^, amide II: 1,555 cm^-1^, ester carbonyl: 1,740 cm^-1^, phosphate: 1,085 cm^-1^). Fig. S5 shows an example of the IR spectra at each step of pre-processing.

For imaging live fibroblast cells, DF images were acquired at 1,502 cm^-1^ and 1,663 cm^-1^. These images were processed similarly to the hyperspectral image cube mentioned above, but without using MNF and baseline subtraction.

The contrast-to-noise-ratio (CNR) is defined as the absorbance difference between a point in the sample (*A*_*s*_) and a point in the background (*A*_*bg*_), divided by the standard deviation of the background (𝜎_*bg*_).

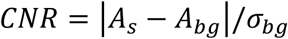

### 2.6 Numerical Simulation

The reflectance spectrum and near field profile of the metasurface sample are simulated using COMSOL Multiphysics (COMSOL, Inc.), a commercially available software using finite elements method. The geometric parameters used were as mentioned in the main text. The nanoantennas are simulated in a unit cell with Floquet boundary conditions in the x and y directions. The optical constants for gold are retrieved from a previous publication.^33^ The structure is simulated with water on the top and supported by CaF_2_ on the bottom, both layers with thickness equal to at least one wavelength. The wave excitation is simulated using a periodic port, where plane waves are incident from the CaF_2_ side.

In order to match the experimentally measured FTIR spectra using the Cassegrain objective, the simulated spectra are calculated as a weighted average of different incident angles with the formula 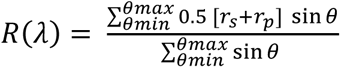, where *θ* is the range of incident angles varying from *θ*_min_ = 10° to *θ*_max_ = 24° with an increment of 1°.^34^ This averaging accounts for the range of angular incidences typical of a 0.4 NA 15x Cassegrain objective. *r*_s_ and *r*_p_ represent the simulated reflectance of incident plane waves in s and p polarizations, respectively.

## 3 Results

### 3.1 Metasurface-enabled MIR imaging

We realized metasurface-enabled DF-MIR chemical using an inverted scanning confocal microscope (see Methods for details). Briefly, the MIR beam from a QCL is focused by an objective (NA = 0.71) onto a diffraction-limited (≈ 5 *μm* full width at half maximum (FWHM) at λ = 6 *μm*) spot, which is laterally scanned by moving a stage-mounted sample comprising the metasurface-bottomed chamber filled with the analyte. The reflected light modulated by the near-field interaction of the analyte with the metasurface is collected by the same objective and measured by a liquid-nitrogen cooled mercury-cadmium-telluride (MCT) MIR detector. The diffraction-limited performance of the imaging optics is validated using a chrome-on-glass 1951 USAF Resolution Test Target to experimentally measure its modulation transfer function (MTF, Fig. S2).^35^

Metasurface-enabled image formation by MIRIAM platform was experimentally investigated by fabricating imaging targets – sub-wavelength polymethyl methacrylate (PMMA) blocks shown in Fig. 2b,e – atop of a single-resonant metasurface (SRM) comprising a periodic array of nanoantennas: see Fig. 2a,d for geometric parameters of the SRM and PMMA blocks. The broad dipolar resonance of the SRM centered at *ɷ*_SRM_ ≈ 1,650 cm^−1^ (in H_2_O) with a linewidth of Δ*ɷ*_SRM_ ≈ 400 cm^−1^ spectrally overlaps with several vibrational bands characteristic of proteins and lipids, including: amide II (*ɷ*_AII_ ≈ 1,550 cm^−1^), amide I (*ɷ*_AI_ ≈ 1,660 cm^−1^), and ester carbonyl 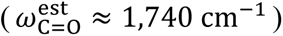 bands. This basic SRM design has identical nanoantennas arranged at sub-wavelength periodicity, smaller than the focal spot size of the IR beam (≈ 5 *μm*). This ensures that multiple nanoantennas are always excited as the sample is scanned through the laser beam, resulting in the reflectance across the analyte-free metasurface being highly uniform in the absence of cells – a requirement for forming high-quality images. See Supplementary Methods 2 and Fig. S3 for details on the geometry and electromagnetic responses of the SRM.

The PMMA targets arrayed with period *P*_3_ = 10*P*_1_ were either centered on (Figs. 2a, b) or between (Figs. 2d, e) the nanoantennas. Hyperspectral data cubes of the reflectance *R*(*x*, *y*; *ɷ*_*j*_) were obtained at each scanned pixel location (*x*, *y*) at DFs *ɷ*_*j*_ of the QCL, from 1,600 cm^-1^ to 1,800 cm^-1^, with 4 cm^-1^ spacing. The *relative* absorbance due to the presence of the target – which can be positive or negative – was defined as *A*(*x*, *y*; *ɷ*_*j*_) ≡ − log_10_(*R*/*R*_0_), where *R*_0_ is the bare SRM reflection in deionized H_2_O. Single-frequency slices of the hyperspectral absorbance cubes, 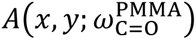, were acquired at the C=O stretching line of PMMA 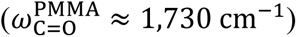 and plotted in Figs. 2c,f for the “on-antennas” (Fig. 2c) and “between-antennas” (Fig. 2f) cases, respectively.

For the “on-antennas” case, each PMMA block is resolved with high contrast, appearing as a Gaussian-shaped spot with FWHM of 5.23 µm (blue line in Fig. 2g). On the other hand, for the “between-antennas” case, each PMMA block is imaged with much weaker contrast and larger spread (FWHM = 33.67 µm: red line in Fig. 2g). Spectral properties of the PMMA obtained by spatial averaging the of the absorbance data (see Methods for details) reveal that the strongest signal is indeed obtained at the frequency corresponding to the C=O stretch.

These results are consistent with the theory that the image contrast is generated primarily by the near-field interaction between the analyte (in this case, PMMA blocks) and the nanoantennas. When the analyte interacts with the metasurface in the near-field of each individual nanoantennas, the analyte can be imaged with diffraction-limited resolution. Note that this would not be the case for imaging systems based on guided wave or distributed feedback photonic elements. For example, imagers based on gratings and photonic crystals, as well as surface plasmon polariton (SPP) based systems, are known to suffer from poor lateral resolution due to the existence of long-range guided modes.^34,36–39^ Such modes appear to be largely absent for the SRM studied here, as revealed by the weak perturbation to the SRM reflection introduced by the PMMA blocks in the “between-antennas” case. This is a somewhat surprising result, since plasmonic nanoantenna arrays are known to support collective surface lattice resonance.^25^ We note that the reflection from PMMA blocks positioned outside of the nanoantenna array was too weak to be seen experimentally. This implies that the finite (albeit low) contrast of the image shown in Fig. 2f can only be explained by the presence of finite radiative coupling between neighboring nanoantennas, but this coupling is small. Therefore, the limited contribution of the radiative nanoantenna coupling of the present SRM – which, for some metasurfaces, can extend over several periods even in the absence of guided modes^40^ – is responsible for diffraction-limited imaging of the MIRIAM platform.

### 3.2 Imaging of fibroblast cells

Multi-spectral imaging of fixed cells using the MIRIAM platform was accomplished using a bi-resonant (BRM) metasurface designed to extend the spectral coverage from the proteins/lipids spectral band to a second spectral band covering the nucleic acids PO_2_^-^ phosphate vibrations. Briefly, the BRM design is based on the self-similar nanoantenna arrays,^41^ and it covers two spectral bands centered around 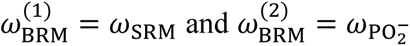, where 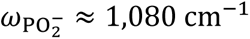. The details of the BRM design are presented in Supplementary Methods 2 and Fig. S3. We have characterized the sensitivity of this BRM to bovine serum albumin (BSA, see Supplementary Methods 3 and Fig. S4) and found that the limit of detection for BSA was remarkably low at LOD_BSA_ = 50 nM, mainly attributed to the high sensitivity of the plasmonic metasurface to surface-adsorbed analyte molecules, confirming the exceptional surface-sensitivity of the plasmonic metasurface.

As a model system for exploring the capabilities of the MIRIAM platform, we have imaged formaldehyde-fixed 3T3-L1 mouse embryo fibroblast cells cultured on the BRM attached to the bottom of a cell culture chamber.^31^ DF-MIR imaging was performed in the two BRM bands: (1) 1,500 – 1,800 cm^-1^ and (2) 1,050 – 1,130 cm^-1^ to form a hyperspectral absorbance cube *A*(*x*, *y*; *ɷ*_*j*_) at Δ*ɷ* ≡ *ɷ*_*j*+1_ − *ɷ*_*j*_ = 5cm^−1^ frequency intervals. Based on the current imaging speed of the point-scanning MIRIAM system, which is about 5 min per wavenumber to image an 800 μm × 600 μm area with 2 μm pixel size, fixation was found to be necessary for hyperspectral imaging because of the cells movement. Faster image acquisition can be expected by scanning the laser beam using galvo scanners ^8^ or by utilizing focal plane arrays (FPA) for wide-field MIR imaging^5,32^, which will be explored in our future work.

In metasurface-enabled chemical imaging, cellular contrast is ultimately determined by the local variation of the complex-valued frequency-dependent dielectric permittivity *ε*(***x***, *ɷ*)/*ε*_0_ ≡ 𝑛^2^(***x***, *ɷ*) of a cell

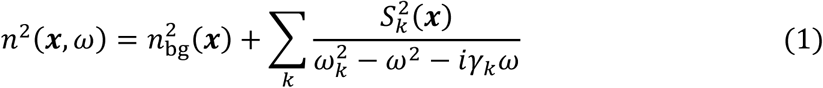

where *ε*_0_is the dielectric permittivity of vacuum, n is the refractive index, n_bg_ is the non-resonant refractive index of the analyte, the summation is over all infrared vibrational modes, ω_k_ is the resonance frequency of the k-th absorption line, γ_k_ is the decay rate, and S_k_ is the oscillator strength. In the MIR spectral range, *ε*(***x***, *ɷ*) has two components: one originates from the absorption and spectral dispersion associated with resonant molecular vibrations (the sum in Equation 1), while the remaining component originates from the non-resonant refractive index variation within the cells (n_bg_). When cells are imaged at a wavenumber *ɷ* = *ɷ*_*j*_ outside of any absorption line, the non-resonant refractive index of the cells 𝑛_bg_(***x***) (which is generally higher than that of culture medium) red-shifts the local nanoantenna resonance at the corresponding location ***x***. This appears as a higher reflectance (and lower relative absorbance *A*(*x*, *y*; *ɷ*_*j*_): see Fig. 3a) on the red-detuned side of the plasmonic resonance, and lower reflectance on the blue-detuned side (not shown). In this way, 𝑛_bg_also produces a baseline absorbance shift: such a shift in the case of PMMA targets can be seen in Fig. 2h. This baseline shift needs to be subtracted to obtain chemical images corresponding to specific vibrational lines. Moreover, the non-uniform 𝑛_bg_(***x***) can be used to visualize the cells through the contrast originating from the local shifts of the plasmonic nanoantenna resonance. Such image collected at *ɷ*_RI_ = 1,500cm^−1^ is shown in Fig. 3a; it can be useful for visualizing different cellular organelles because of their distinct refractive index. For example, cellular nuclei are clearly visible in Fig. 3a because of the higher refractive index within the nucleus.

**Fig. 3.**
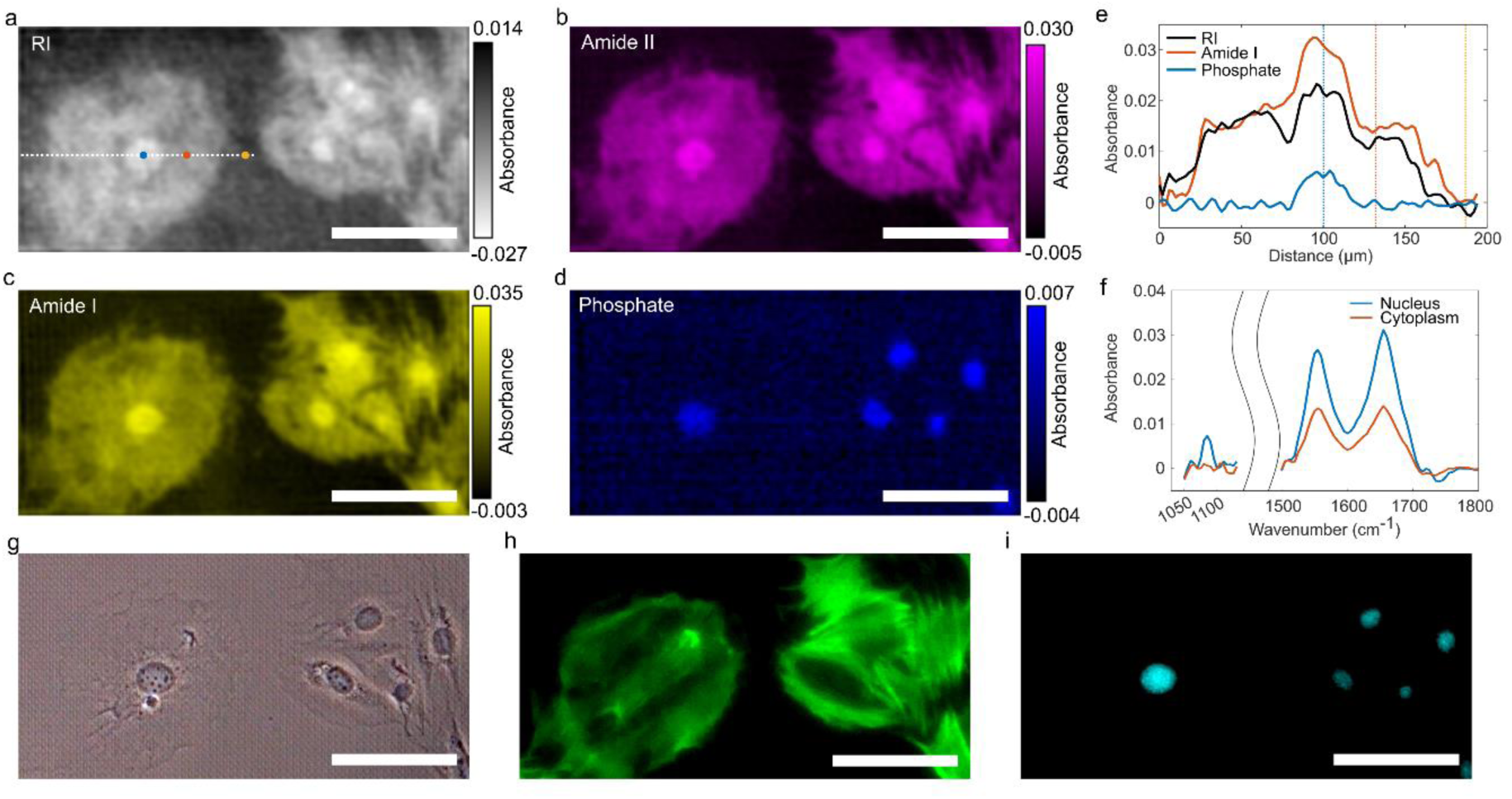
Multi-spectral imaging of fixed 3T3-L1 fibroblast cells using the MIRIAM platform. (a-d) Chemical images of the selected region of the metasurface with adhered cells collected at (a) no molecular vibration 𝝎_𝐑𝐈_ = **1**, 𝟓𝟎𝟎 𝐜𝐦^−**1**^, (b) amide II (𝝎_𝐀𝐈𝐈_ ≈ **1**, 𝟓𝟓𝟓 𝐜𝐦^−**1**^), (c) amide I (𝝎_𝐀𝐈_ ≈ **1**, 𝟔𝟓𝟓 𝐜𝐦^−**1**^), and (d) PO_2_^−^ phosphate 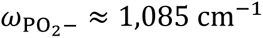. Blue, red, and yellow dots in (a) mark the position of the nucleus, cytoplasm, and background, respectively. Blue, red, and yellow dotted lines mark the different positions, corresponding to the dots in (a). (e) Line profile of the absorbance signal along the white dotted line marked in (a), for different vibrational contrasts. (f) MIR absorbance spectra inside the nucleus (blue line) and cytoplasm (red line) at the locations indicated in (a). (g-i) Phase contrast image (g), actin stained (h), and nucleus (DAPI) stained (i) fluorescence microscopy images of the same cells. Scale bars: 100 µm.

The absorbance images, corrected by the linear baseline subtraction for the above-mentioned non-vibrational absorbance shift, are plotted in Figs. 3b-d for several frequencies (amide II at *ɷ*_AII_ ≈ 1,555 cm^−1^, amide I at *ɷ*_AI_ ≈ 1,655 cm^−1^, and PO_2_^-^ phosphate at *ɷ*_PO2−_ ≈ 1,085 cm^−1^) to reveal the corresponding vibrational contrast images. As expected, the two amide absorbance images shown in Figs. 3b-c are very similar because both correspond to the protein-related contrast. Both images clearly show the overall morphology of the cells and sufficiently high contrast-to-noise ratio (CNR ∼ 40) for clear observation of the cell boundary.

For comparison, the corresponding phase contrast image (Fig. 3g) reveals weak contrast for these fibroblasts, most likely because of their thin/flat morphology. When compared to the actin-stained fluorescence image of the same cells (Fig. 3h), the MIR images reveal fiber-like features attributed to filopodial actin and stress fibers. We also note that the MIR images exhibit higher contrast uniformity within each cell than the corresponding fluorescence image, suggesting significant contribution to the absorbance from proteins other than actin.

Just as in the non-resonant refractive index (RI) image in Fig. 3a, the cellular nuclei produce stronger protein contrast than the cytoplasm because of the higher protein density inside the nuclei. This observation agrees with a previous study based on MIR photothermal microscopy.^15^ Moreover, the MIRIAM platform resolves the unique chemical compositions of different cellular organelles. For example, Figs. 3b-d reveal co-localization of the nucleic acid-associated phosphate PO_2_^−^ (Fig. 3d) and protein-associated (Figs. 3b,c) amide I/II absorbance within the cell nuclei, but not in the rest of the cell. This is clearly visualized from the absorbance line profile across the cell from different vibrational contrast (Fig. 3e). Comparison between the DAPI-stained fluorescent (Fig. 3i) and the PO_2_^−^ band MIR images confirms that the latter is indeed associated with the nuclei. The spectra inside the nucleus/cytoplasm (Fig. 3f), extracted from the hyperspectral absorbance cube at the locations marked by the blue/red dots in Fig. 3a, clearly show the difference between the two organelles: the nucleus is characterized by high absorbance at the amides and phosphate bands, whereas the cytoplasm exhibits weaker absorbance at the amide bands and essentially no signal at the phosphate band.

Given that the MIRIAM technique is most sensitive to analytes within the “hotspots” at the nanoantenna tips, and considering that the penetration depth of these hotspots is on the order of 100 nm, it is not surprising that the absorbance attributed to membrane proteins, focal adhesion complexes, and cytoskeletons can be seen. The high contrast of the nuclei-associated images is not immediately intuitive because of their deeper location inside the cells. We note, however, that the fibroblast cells are known to have a spread-out flat morphology.^42,43^ The spreading of the cell body likely brings the nucleus closer to its bottom, within the penetration depth of the nanoantenna near-field.

### 3.2 Imaging of adipocyte cells

Next, we have induced adipocyte differentiation in 3T3-L1 cells and allowed the lipid droplets (LDs) to mature, to investigate the metasurface-enabled MIR imaging of LDs inside the adipocytes. LDs are dynamic cellular organelles that serve as storage sites for neutral lipids.^44^ Because of their critical role in lipid metabolism, the study of LDs using vibrational microscopy has been an active area of research.^15,45,46^ Although the result for fixed cells are presented here due to our limited acquisition rate, live adipocyte imaging were performed too by imaging at 5 DFs (Fig. S6), and the images at different vibrational bands look very similar to the fixed cell images.

MIR absorbance images at *ɷ*_AII_ and *ɷ*_AI_frequencies (Figs. 4a and 4b, respectively) show that individual cells are resolved, and the cell morphologies are clearly seen from the protein-derived contrast. Additionally, we observe small dark circular objects corresponding to negative absorbance in the amide I image shown in Fig. 4b. These objects are co-located with LDs visible in the BF image shown in Fig. 4d. Similar dark contrast in LDs has also been observed previously with MIR optoacoustic microscopy.^46^ A plausible interpretation of the negative absorbance of LDs is that they have completely displaced H_2_O from their volume. Because H_2_O has a strongly-absorbing OH bending vibrational mode at *ɷ*_H−O−H_ ∼ 1,650 cm^−1^, spectrally overlapped with the amide I band, its displacement manifests as a negative relative absorbance. This effect is clear from the MIR absorbance spectra (Fig. 4e) collected at the two cellular locations labelled in Fig. 4b: inside (red arrow) and outside (blue arrow) of the LD. The absorbance inside the LD (red line in Fig. 4f) is clearly diminished around *ɷ*_H−O−H_ ∼ 1,650 cm^−1^ when compared with that outside the LD (blue line) because of the suppressed H_2_O absorbance. Nevertheless, both amide I and II absorbance bands are still clearly expressed inside the LD spectrum; those are attributed to the cytoskeletal and membrane proteins located below the LD but above the metasurface. The difference spectrum between the inside and outside LD locations (yellow dashed line in Fig. 4e) clearly shows the water-associated absorption dip (negative absorbance) around *ɷ*_H−O−H_ ∼ 1,650 cm^−1^, and the prominent C=O vibrational peak at 1,740 cm^-1^ can be clearly seen. This difference likely better represents the absorbance of the LD alone, without contribution from the proteins below.

**Fig. 4.**
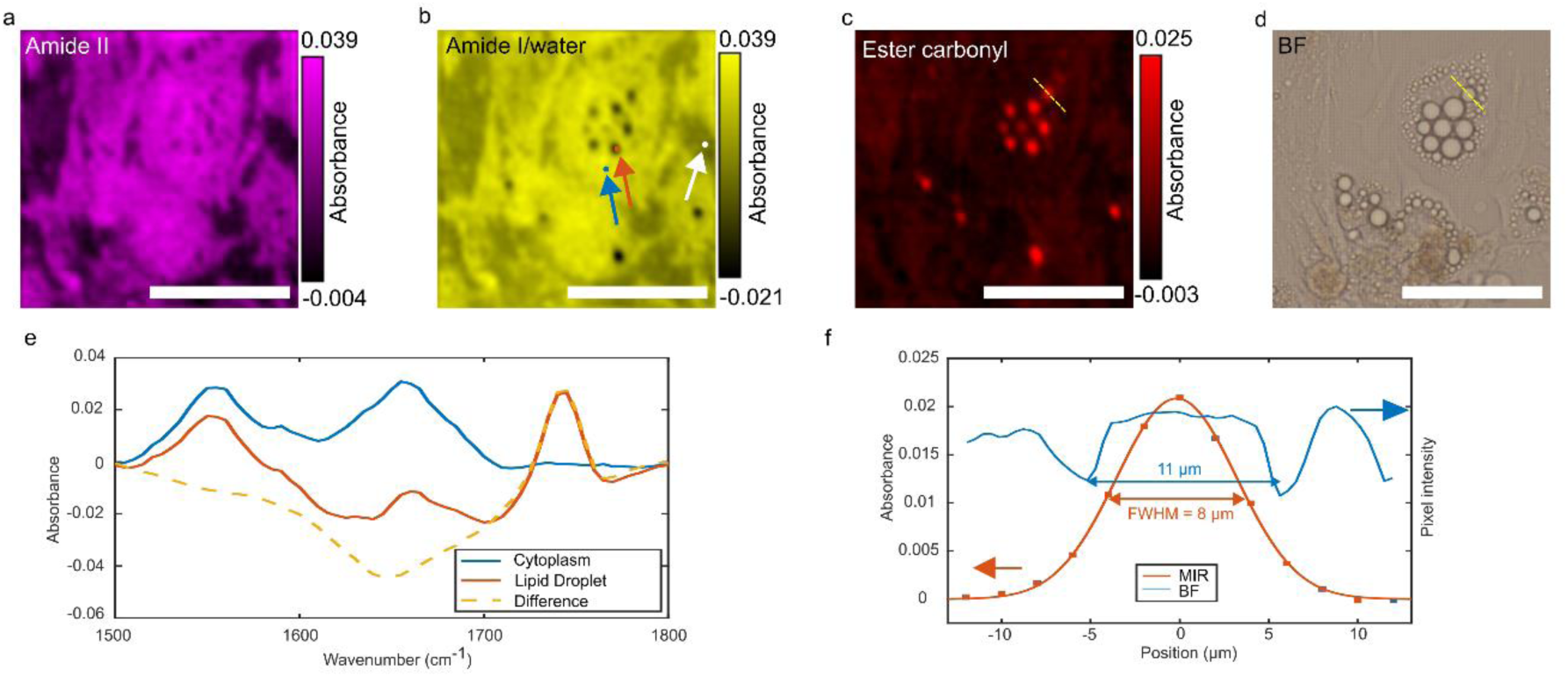
Multi-spectral imaging of fixed 3T3-L1 adipocytes using the MIRIAM platform. (a-d) Chemical images of the selected regions of the metasurface with adhered cells collected at (a) amide II (𝝎_𝐀𝐈𝐈_ ≈ **1**, 𝟓𝟓𝟓 𝐜𝐦^−**1**^), (b) amide I (𝝎_𝐀𝐈_ ≈ **1**, 𝟔𝟓𝟓 𝐜𝐦^−**1**^), and (c) ester carbonyl 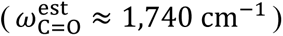 frequencies. LDs appear as dark spots in (b) because of the overlapping OH bending absorption band from the displaced water. Blue, red and white arrow/dots mark the position of cytoplasm, lipid droplet, and background, respectively. (d) Brightfield (BF) image of the same cells. (e) Relative absorbance spectra inside the cytoplasm (blue line) and LDs (red line) at the locations marked in (b). Yellow line: the difference between the cytoplasm and LD spectra. (f) Intensity profile along the yellow dashed line in (c) and (d). Red dots are the absorbance intensity along the yellow dashed line in (c), and the red curve is a Gaussian fit. Blue curve represents the normalized grayscale pixel intensity along the yellow dashed line in (d). Scale bars: 100 µm.

While the composition of the LDs can only be conjectured from the BF image, the MIR imaging platform enables definitive determination of their chemical composition. The lipid composition of these small circular organelles is validated by the MIR absorbance image at 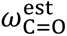 (see Fig. 4c) corresponding to the ester carbonyl C=O band. Individual organelles observed in the lipids-associated image in Fig. 4c are co-located with the negative absorbance objects shown in Fig. 4b, thus validating their interpretation as LDs. By comparing the MIR and BF images, we note that only large LDs (diameter > 10 μm: see Fig. 4d) are observed in Figs. 4b,c; the smaller LDs are entirely absent from the MIR image. In addition, the LDs in the MIR images appear to be smaller than their counterparts in the BF image. For instance, the LD indicated by the yellow dashed line in Fig. 4c,d has a FWHM ∼ 8 μm size from the MIR image, but appears to have a larger (FWHM ∼ 11 μm) size according to the BF image: see Fig. 4f. These differences between the BF and MIR images are most likely related to the shallow penetration depth of the metasurface nearfield. Due to its lipid core, LD is more buoyant than most cellular organelles.^47^ We conjecture that smaller LDs float to the top of the cell, placing them too far away from the metasurface to be observable by the MIRIAM platform. On the other hand, the larger LDs could be spatially confined by the finite thickness of the cell and pushed toward the metasurface and into its nearfield. However, even for large LDs, only their bottom portion overlaps with the evanescent optical field of the metasurface. This could explain the apparently smaller size of the LDs in the MIR image.

### 3.3 Time-lapse imaging of living cells

Finally, we demonstrate the ability of the MIRIAM platform to perform time-lapse imaging of living cells undergoing adhesion/detachment, spreading, and locomotion. Based on the time scale of these processes, we have established the acceptable acquisition rate (Δ𝑡 =7 minutes per frame) and the number of frequencies (two: *ɷ*_RI_ = 1,502 𝑐𝑚^−1^ and *ɷ*_AI_ = 1,663 𝑐𝑚^−1^) compatible with such time resolution under the constraints of our current image acquisition speed. The resulting MIR images capture the overall cellular RI contrast (Fig. 5) and the protein contrast derived from amide I absorption (Fig. S7). Note that no baseline subtraction was applied to form these images, thus although the amide I band image is still dominated by protein contrast, it is not pure, and may have some refractive index contrast from plasmonic resonance shift mixed in. All 60 frames of these two time-lapse videos, for the total time of 𝑇_tot_ ≈ 410 min, are included in the supplementary materials.

**Fig. 5.**
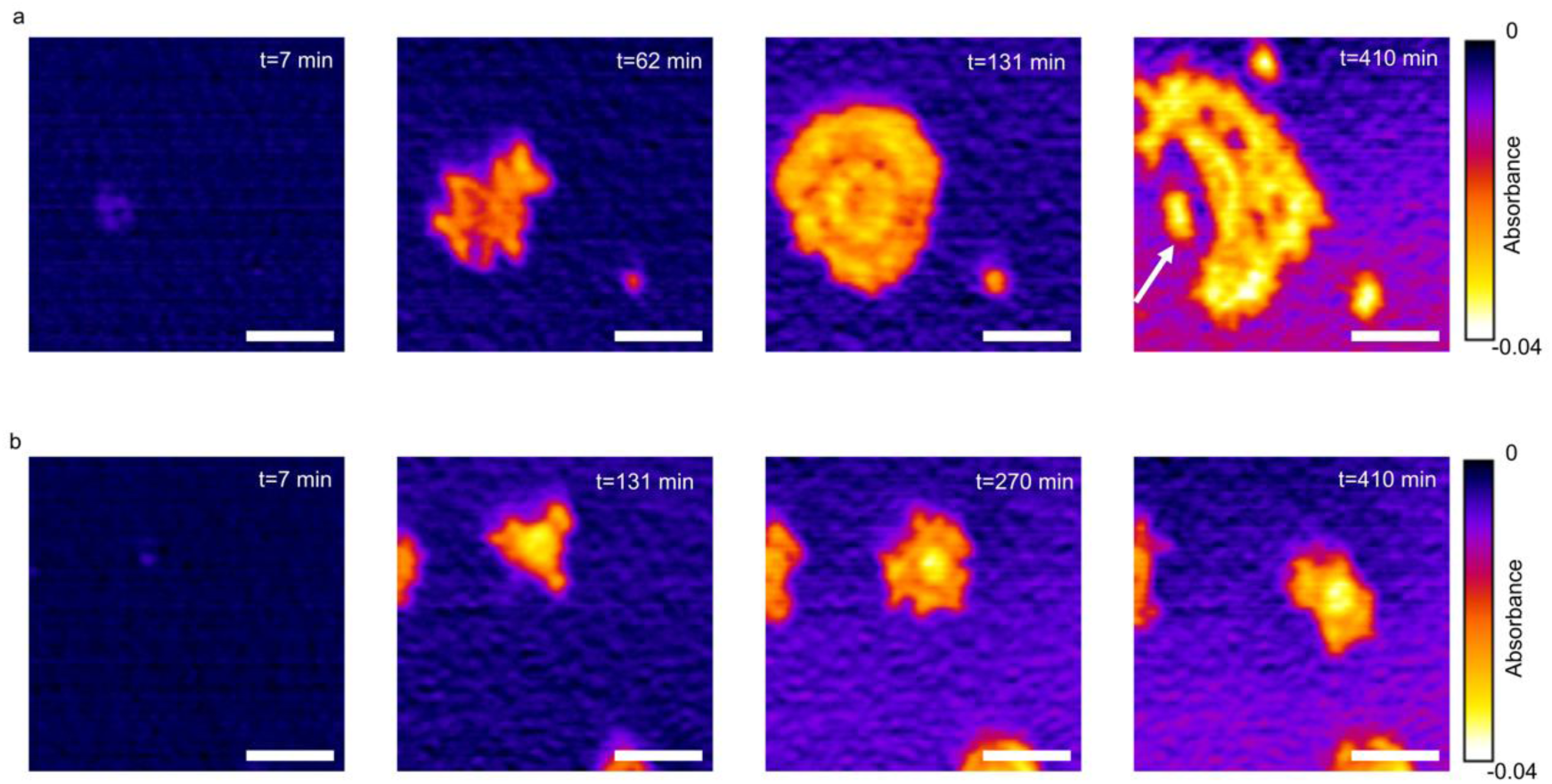
Time-lapse imaging of living 3T3-L1 fibroblasts. (a) Adhesion and spreading of a fibroblast cell on the metasurface over 7 hours. (b) Locomotion of a fibroblast cell on the single-resonant metasurface (SRM). Contrast: relative absorbance at 𝝎_𝐑𝐈_ = **1**, 𝟓𝟎𝟎 𝐜𝐦^−**1**^corresponding to local RI-induced shift of the MIR plasmonic resonance of the nano-antennas. Scale bars: 50 µm.

After 3T3-L1 fibroblast cells were seeded on the metasurface, the time-dependent absorbance hypercube *A*(*x*, *y*, 𝑡; *ɷ*_*j*_) was measured during the course of their adhesion, spreading, and locomotion on the SRM. Fig. 5 (a) presents an example of a cell that quickly initiates adhesion to the metasurface, then spreads over a large area over the course of the imaged period. The signal strength is highly non-uniform within a single cell, likely mirroring the distribution of cellular focal adhesion. Notably, several “hollows” are observed within the cell at t = 410 min, and a portion of the cell (white arrow) appears detached from its main body. At these positions the cell body is presumed to be too far above from the metasurface to be imaged in the near-field of the metasurface, highlighting the surface-sensitive nature of the MIRIAM imaging platform.

Moreover, this example underscores the importance of recording the entire time-lapse video, in the absence of which the above-mentioned portion of the cell could have been misinterpreted as a separate small cell. For example, some of the cells (see Fig. 5b) remained fairly small over the same time period. Note that the cells in Fig. 5b are quite motile and exhibit clear locomotion. In both cases, continuous changes in cellular morphology and the formation and dissolution of filopodia around the cell periphery could be observed with MIRIAM using a single QCL frequency. Time lapse images with larger field-of-view, including both regions of interest presented in Fig. 5, is included in Fig. S8.

These time-lapse imaging results clearly demonstrate that living cells can be imaged using the MIRIAM platform, thereby paving the way for studying diverse dynamic cellular processes through vibrational imaging. Furthermore, these results indicate that cells can remain viable over an extended period (7 hours or more) during imaging, and that the MIR light from QCL does not induce significant heating that impacts cellular behavior and viability.

### 3.4 Other cell lines

To demonstrate that MIRIAM is applicable to a variety of cell lines beyond 3T3-L1, we performed imaging of three human-derived cancer cell lines with protein contrast (amide II): A431 human epidermoid carcinoma cells, GUMC387 conditionally-reprogrammed human prostate cancer cells, and MDA-MB231 human breast cancer cells (Fig. S9). The distinct morphologies of these cell lines are clearly visible in the protein contrast images. These results highlight the broad applicability of MIRIAM as a cellular imaging technique.

## 4 Discussion

In this work, we have demonstrated metasurface-enabled MIR chemical imaging, where the cells on a plasmonic metasurface are imaged through their near-field interactions with the nanoantennas of the metasurface. This approach offers non-destructive, label-free imaging of live cells, providing vibrational contrast for various cellular organelles, including the nucleus, cytoplasmic proteins, and LDs. With diffraction-limited resolution in the MIR, we achieve a spatial resolution of approximately 5 μm, enabling visualization of relatively large cellular organelles such as nuclei and LDs.

Our reported technique introduces a unique imaging approach, where images are formed by sampling the analyte at periodic plasmonic hotspots of the MIR field in the immediate proximity of the nanoantennas, with the analyte located within these hotspots predominantly contributing to the signal. This stands in contrast to typical imaging methods, where all analytes within the probe beam focal spot contribute to the signal. When designing the metasurfaces for the MIRIAM platform, we became aware of the tradeoff between utilizing surface lattice resonances resulting from far-field coupling between nanoantennas (which can lead to higher quality factor of the plasmonic resonance and facilitate imaging of analytes outside the plasmonic hotspot) and avoiding such far-field coupling (which improves lateral spatial resolution). This differs from the design principle of conventional plasmonic nanoparticle or nanoantenna-based biosensors, where surface-lattice resonance is viewed as entirely beneficial due to its role in narrowing the plasmonic resonance linewidth. As further investigations into metasurface-based imaging are pursued, additional research on optimal metasurface design would be valuable and could potentially achieve spatial resolution beyond the diffraction limit, ultimately limited by the periodicity of the nano-antennas.

Compared to existing MIR imaging techniques, the MIRIAM approach stands out for its exceptional surface sensitivity. This is evidenced by its high sensitivity to adsorbed proteins like BSA, as well as its ability to provide high contrast for imaging cell morphology and adhesion on the surface. This surface sensitivity, however, does present challenges when imaging structures deeper inside the cells, although imaging of cell nuclei and LDs remains feasible. Integrating patterned nano-topographies and nanopillars with plasmonic nanoantennas could be one way to address this issue.^48^

The plasmonic metasurface used in this work is fabricated on a planar substrate and is fully compatible with standard cell culture vessels, such as petri dishes and microwell plates. There is no need to confine the cell within a thin culture chamber, thus enhancing the long-term viability of the cells. Additionally, there are no extra optics or detectors above the metasurface, potentially making the metasurface device compatible with modern liquid and plate handlers. These advantages make our proposed technique especially suitable for the MIR hyperspectral imaging of living cells in multi-well plate format, with potential applications in high-throughput drug screening. In particular, MIR vibrational probes can be employed to track distinct metabolic activities, making the high-throughput metabolic phenotyping of living cells a highly promising application of our proposed technique. While our metasurface is currently fabricated using electron-beam lithography, wafer-scale fabrication based on deep-ultraviolet (DUV) lithography can be employed to mass-produce metasurfaces for such applications.^49^

The slow image acquisition rate poses a limitation in our current setup, restricting us from imaging at more than 2-3 wavenumbers for living cells that exhibit dynamic changes in morphology. In our setup, the acquisition speed was limited by the scanning speed of the microscope stage. Significant progress has been made in mid-infrared (MIR) microscopy in recent years, and the development of MIR dual-axis galvo laser scanning microscopes with corrected optics for the MIR range has been reported, greatly accelerating image acquisition.^8^ Alternatively, wide-field MIR imaging based on microbolometer or MCT focal plane array (FPA) can also facilitate faster image acquisition. Our metasurface is compatible with either of these imaging techniques, thus the reported technique can be seamlessly integrated with state-of-the-art MIR microscopes for the high-speed chemical imaging.

## Code, Data, and Materials Availability

Code, data, and materials are available upon request from the corresponding author.

## Acknowledgments

The research reported here was supported by the National Cancer Institute of the National Institutes of Health under award number R21 CA251052 and by the National Institute of General Medical Sciences of the National Institutes of Health under award number R21 GM138947 (to G. Shvets). This work was performed in part at the Cornell NanoScale Facility, a member of the National Nanotechnology Coordinated Infrastructure (NNCI), which is supported by the National Science Foundation (Grant NNCI-2025233).

## Author contributions

S.H.H., P.-T.S., and G. Shvets designed the experiment. S.H.H., P.-T.S., and G. Sartorello built the optical setup. S.H.H. and P.-T.S. performed the experiment and analyzed the data. A. M. performed the numerical simulation. J.L. and X.L. provided cell line samples for the experiment. G. Shvets supervised the project. S.H.H. and P.-T.S. wrote the paper with revision by G. Shvets and contribution from all authors.

## Supplementary Text

### Performance of the MIR imaging system

We characterized the imaging capabilities of our MIR laser confocal microscope setup using a chrome-on-glass 1951 USAF Resolution Test Target (Fig. S2). The reflectance image shows that patterns in group 6 are clearly resolved when imaged at λ = 6 µm (1,667 cm^-1^). The modulation transfer function (MTF) was obtained experimentally from the reflectance image line profilers of the repeating line bars of the resolution test target. The imaging system resolved element 6-4 (feature size 5.52 µm) of the resolution target with 30% modulation (corresponding to Rayleigh criterion), demonstrating that our imaging system is performing at close to diffraction limit (5.15 µm resolution with NA = 0.71 objective).

### Details of the single-resonant (SRM) and bi-resonant (BRM) metasurfaces

Two metasurface geometries were used in this work: single-(SRM) and bi-resonant (BRM) metasurfaces. The SRM comprises a periodic array of nanoantennas supporting dipolar resonances centered at ω_SRM_ ≈ 1,650 cm^−1^. To extend the spectral coverage to the PO_2_^-^ phosphate vibration 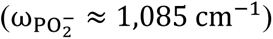 characteristic of the nucleic acids, a BRM design was used. Electromagnetic simulations (Fig. S3b) show that such a BRM supports resonant near-field distribution localized to each set of nanoantennas for each spectral band of interest. In both metasurface designs, the resonant modes only couple to incident light polarization along the long axis of the nanoantennas, and the polarization of the linearly polarized QCL beam was chosen along that direction.

### Characterization of the sensitivity of BRM-based MIRIAM platform to protein and phosphate ions in water

We characterized the sensitivity of the BRM-based MIRIAM platform to bovine serum albumin (BSA) and sodium phosphate dibasic in H_2_O. The relative absorbance *A*(*x*, *y*; *ɷ*_𝑖_) ≡ − log_10_(*R*/*R*_0_) of the analyte at each pixel location (*x*, *y*) in the central part of the metasurface was calculated from the reflectance *R*(*x*, *y*; *ɷ*_𝑖_) normalized to the analyte-free reflectance *R*_0_(*ɷ*_𝑖_) in deionized water.

For BSA, the spatially-averaged absorbance *A*(*ɷ*_𝑖_) ≡ 〈*A*(*x*, *y*; *ɷ*_𝑖_)〉 was calculated from three laser frequencies within the amide I band: ω_1_ = 1,601 cm^-1^ and ω_2_ = 1,702 cm^-1^ for linear baseline subtraction to remove the plasmonic resonance shift, and *ɷ*_*A*𝐼_ = 1,658 𝑐𝑚^−1^ for amide I absorbance (Fig. S4a). Using a range of bovine serum albumin (BSA) concentrations (from 10 nM to 10 µM: see Fig. S4 b), we have determined the limit of detection (LOD) for the BSA solution: LOD_BSA_ ≈50 nM (3.3 µg/mL), which was defined as the lowest concentration with a SNR of 3.

For sodium phosphate dibasic, the spatially averaged absorbance *A*(*ɷ*_𝑖_) ≡ 〈*A*(*x*, *y*; *ɷ*_𝑖_)〉 was similarly calculated with linear baseline subtraction, using absorbance signal at ω_phosphate_ = 1,080 cm^-1^ and using ω_3_ = 1,035 cm^-1^ and ω_4_ = 1,145 cm^-1^ to compute the linear baseline (Fig. S4c). The LOD for sodium phosphate dibasic was determined to be 10 mM (1 mg/mL of monohydrogen phosphate ions), as shown in Fig. S4d.

We note that the sensitivity of the MIRIAM technique to BSA is surprisingly high compared with other vibrational spectroscopy techniques that do not utilize plasmonic metasurfaces: SRS,^19^ MIR photothermal microscopy,^14^ and MIR optoacoustic microscopy^46^ typically have ∼mM LODs for molecular vibrations from small biomolecules in water or D_2_O, and ∼ µM range LODs in case of proteins. The high sensitivity of our technique for BSA is attributed to the surface adsorption of BSA molecules to the gold nanoantennas. The highly nonlinear dependence of the absorbance on the BSA concentration shown in Fig. S4b – the signal is linear and highly sensitive at low BSA concentration, but saturates at higher concentrations – strongly suggests surface adsorption effects.^50^ Such high sensitivity to surface-bound biomolecules highlights the working principle of the MIRIAM as a surface-sensitive technique, and is consistent with previous reports on the detection of protein monolayers using resonant SEIRA metasurfaces.^24,25^ On the other hand, for sodium phosphate, the sensitivity measured from MIRIAM is comparable to other vibrational imaging techniques, and the response is linear, as expected from an analyte without significant surface adsorption.

In addition to its high sensitivity, MIRIAM requires very low optical power (*P*_𝑄𝐶𝐿_ < 14𝜇𝑊) because of its linear nature: it measures the reflected MIR light directly, rather than through the often-weak indirect measurement of either heat or ultrasound generated by a MIR pump, as done in MIR photothermal and optoacoustic microscopy.^14,46^ This not only simplifies the experimental setup, but it also lowers phototoxicity concerns by utilizing MIR laser power that is at least an order of magnitude smaller than the above-mentioned techniques.

**Figure S1.**
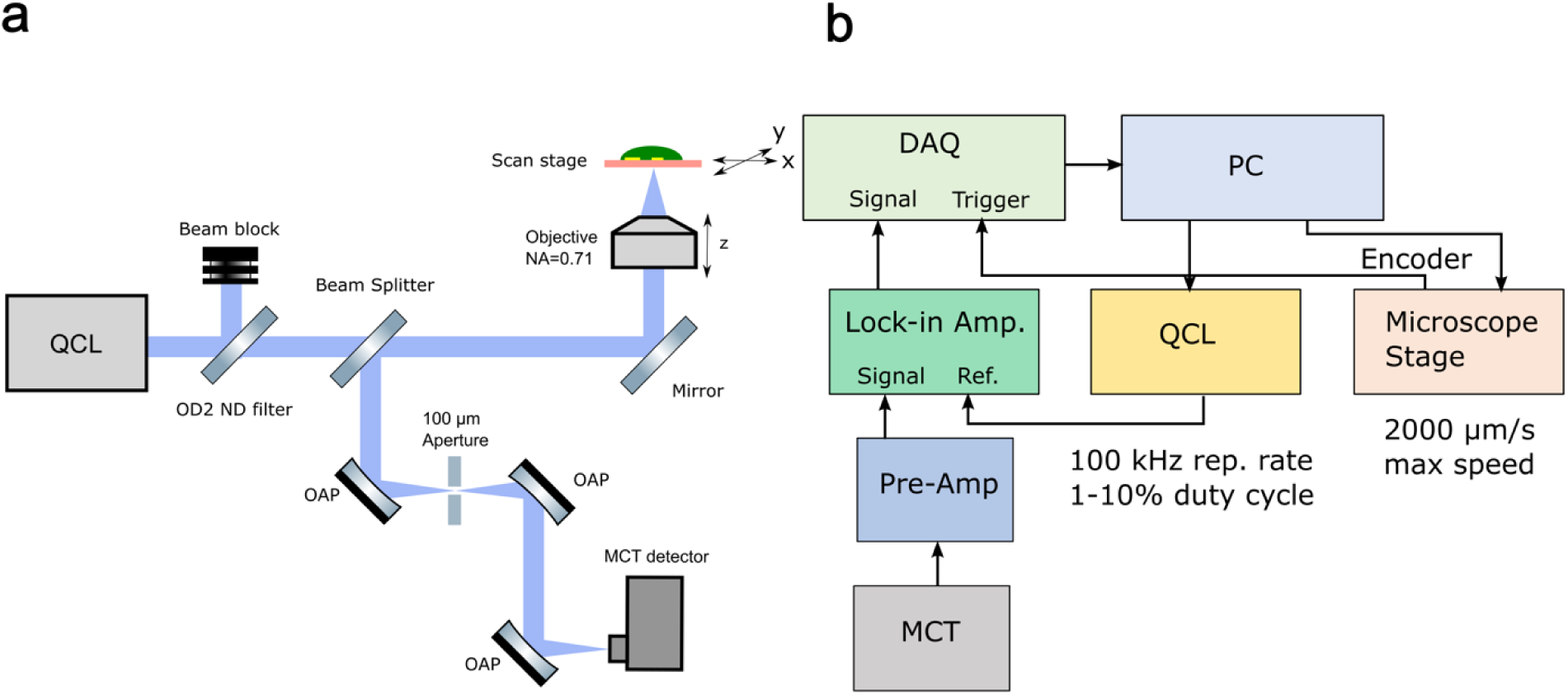
The schematics of the inverted mid-infrared (MIR) microscope, used to image cells on metasurface. (a) The optical setup. OAP, off-axis parabolic mirror; QCL, quantum cascade laser; MCT, mercury-cadmium-telluride; OD2 ND filter, neutral density filter with optical density of 2. (b) The electronic schematics of the system. DAQ, data acquisition card; PC, personal computer.

**Figure S2.**
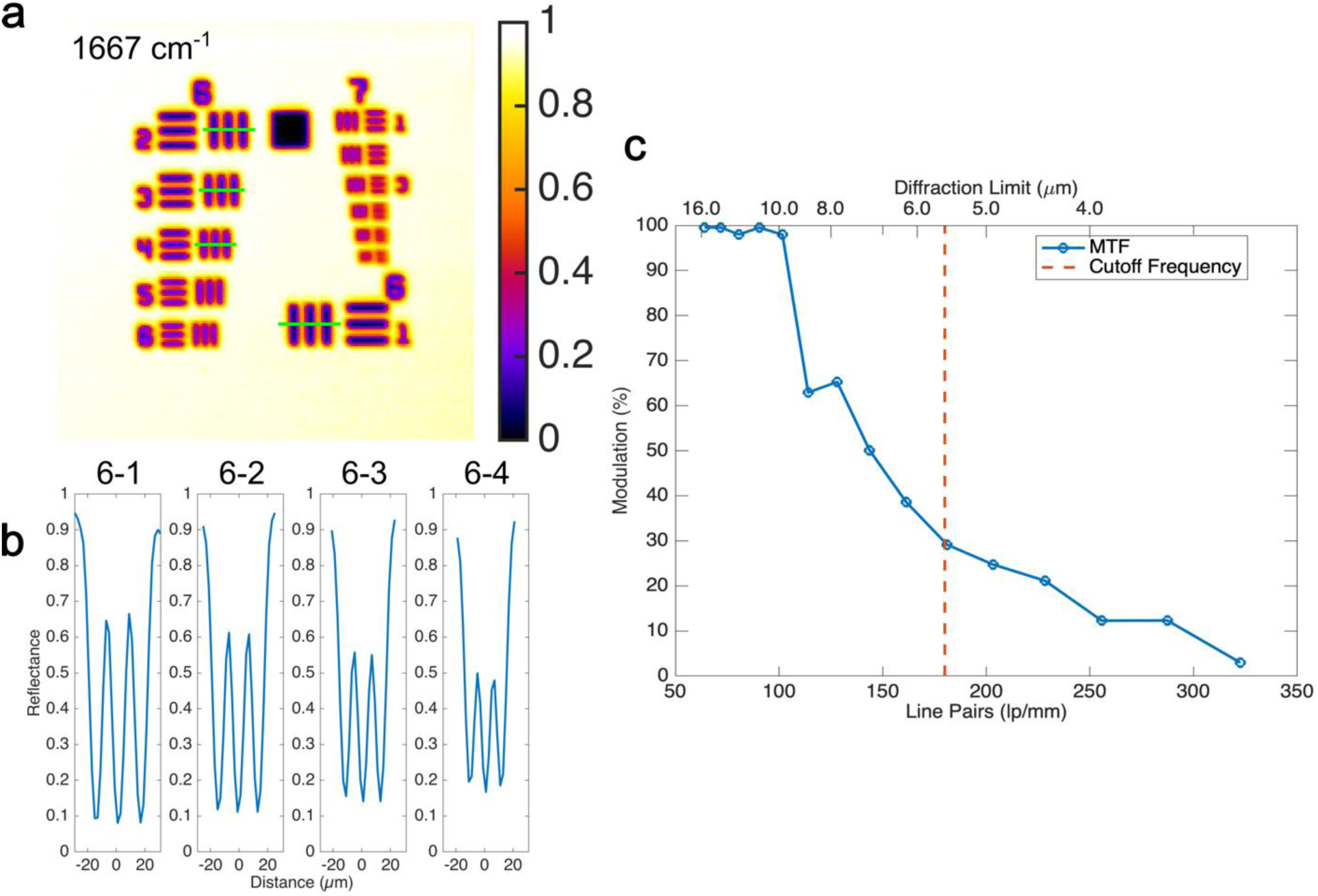
The imaging performance of the optical system. Spatial resolution is demonstrated with a USAF 1951 target (R1L1S1N, Thorlabs). (a) Reflectance image from a USAF 1951 target at 1,667 cm^-1^. The green lines mark the selected line profiles (elements 6-1, 6-2, 6-3, and 6-4) used for modulation transfer function (MTF) evaluation. (b) The line profiles of elements 6-1, 6-2, 6-3, and 6-4. (c) The MTF of the system. 30% modulation was chosen to be the criteria for resolvability, corresponding to 180 lp/mm, which is equivalent to a resolution of 5.56 µm.

**Figure S3.**
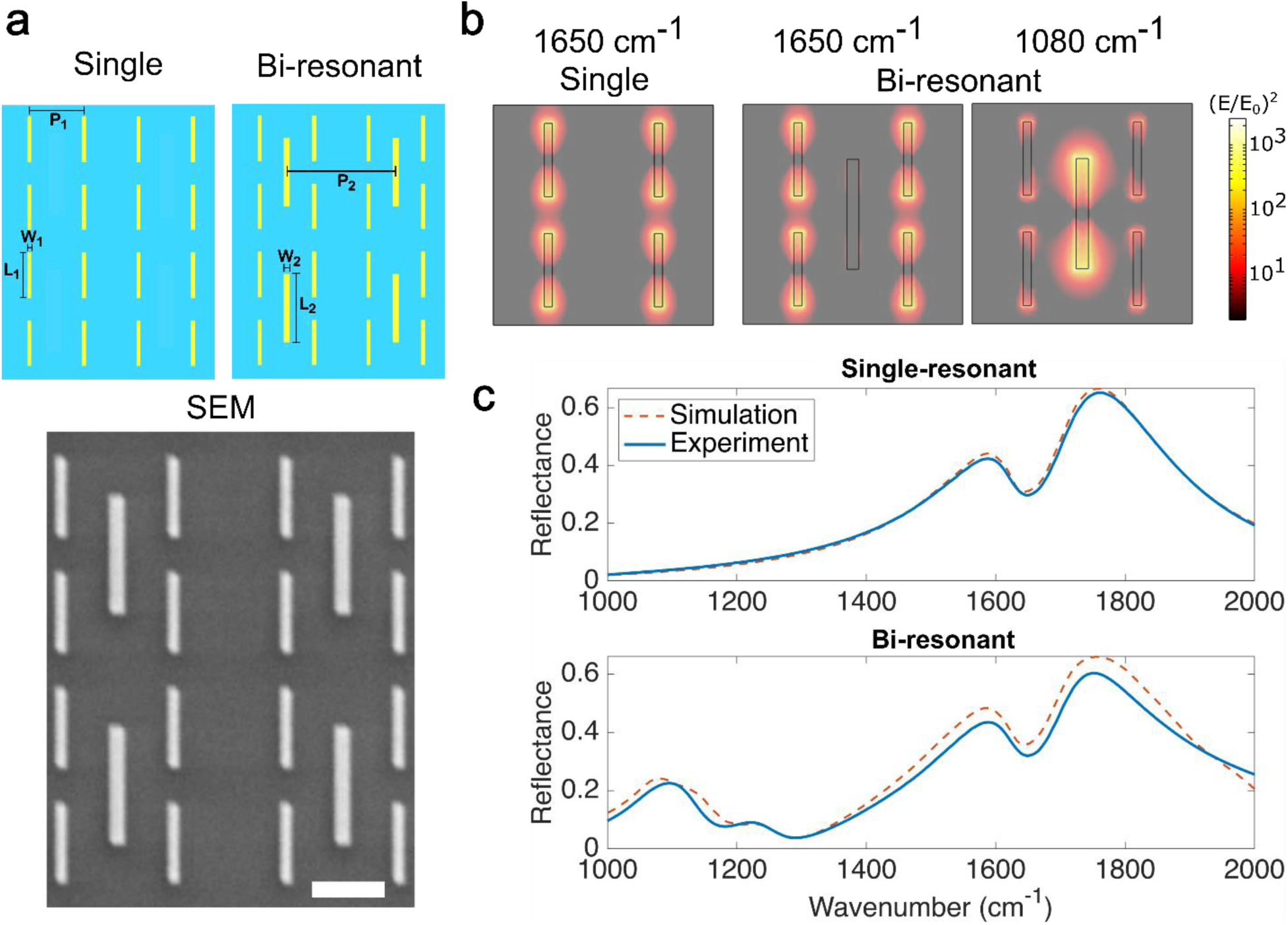
Metasurface design for cell imaging. (a) Top: Schematic drawings of the single and bi-resonant metasurface. The single-resonant metasurface consisted of one type of nanoantenna. The nanoantenna dimension is 1.8 µm × 200 nm with the resonance centered at 1,650 cm^-1^ and a period P_1_ of 2.7 µm. The bi-resonant metasurface consisted of two nanoantennas: a sub-array of the same nanoantennas as the single-resonant metasurface, and an additional sub-array of longer nanoantennas. The longer nanoantenna has a dimension of 2.7 µm × 300 nm with the resonance centered at 1,100 cm^-1^ and a period P_2_ of 5.4 µm. Bottom: Scanning electron microscope (SEM) image of the fabricated bi-resonant metasurface. Scale bar: 2 µm. (b) Simulated electric field intensity profile of the metasurfaces at 1,650 cm^-1^ and 1,080 cm^-1^. The hot spots from dipolar resonance of the nanoantennas can be seen. (c) Experimentally measured reflectance spectra of the single and bi-resonant metasurfaces (blue curve), compared with the simulated spectra (red, dotted curve).

**Figure S4.**
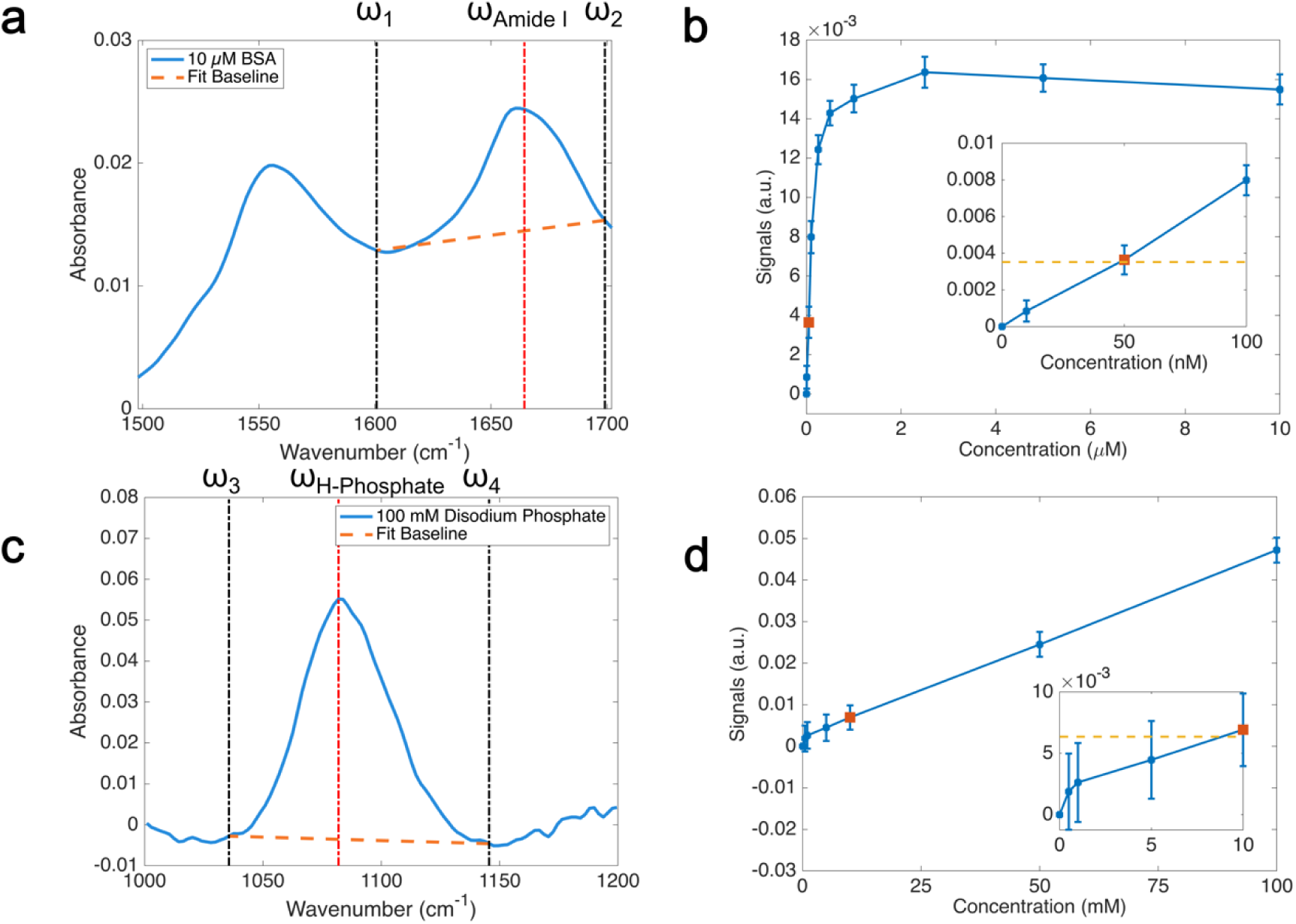
Sensitivity of metasurface-enhanced infrared reflection chemical imaging, for bovine serum albumin (BSA) and sodium phosphate dibasic. (a) FTIR absorbance spectrum of 10 µM BSA on metasurface. A clear sloped baseline can be seen, due to the plasmonic resonance shift. To correct for this baseline, we linearly interpolated the baseline using ω_1_ = 1,600 cm^-1^ and ω_2_ = 1,700 cm^-1^, to obtain the baseline corrected absorbance signal at ω_Amide I_ = 1,660 cm^-1^. (b) QCL-based absorbance signals at varying concentrations of BSA. Absorbance signal was measured at 1,601, 1,658, and 1,702 cm^-1^, slightly offset from the wavenumbers used in the FTIR measurement, to avoid water vapor absorption lines. Error bars represent the standard deviation of the signal. The orange dashed line represents 3 times the standard deviation of signal from water, which defines the criteria for the limit of detection. The highly non-linear response curve is attributed to the adsorption of BSA onto the gold nanoantennas. The limit of detection was 50 nM, or 3.3 µg/mL. (c) FTIR absorbance spectrum of 100 mM sodium phosphate dibasic. Linear interpolation using ω_3_ = 1,035 cm^-1^ and ω_4_ = 1,145 cm^-1^ is used for baseline subtraction, with the baseline-corrected signal obtained at ω_phosphate_ = 1,080 cm^-1^. (d) QCL-based absorbance signals at varying concentrations of sodium phosphate dibasic. Absorbance signal was measured at 1,035, 1,080, and 1,145 cm^-1^. Error bars represent the standard deviation of the signal. The orange dashed line represents 3 times the standard deviation of signal from water, which defines the criteria for the limit of detection. A linear response curve was seen, in contrast to BSA. The limit of detection was 10 mM, or 1 mg/mL of monohydrogen phosphate ions.

**Figure S5.**
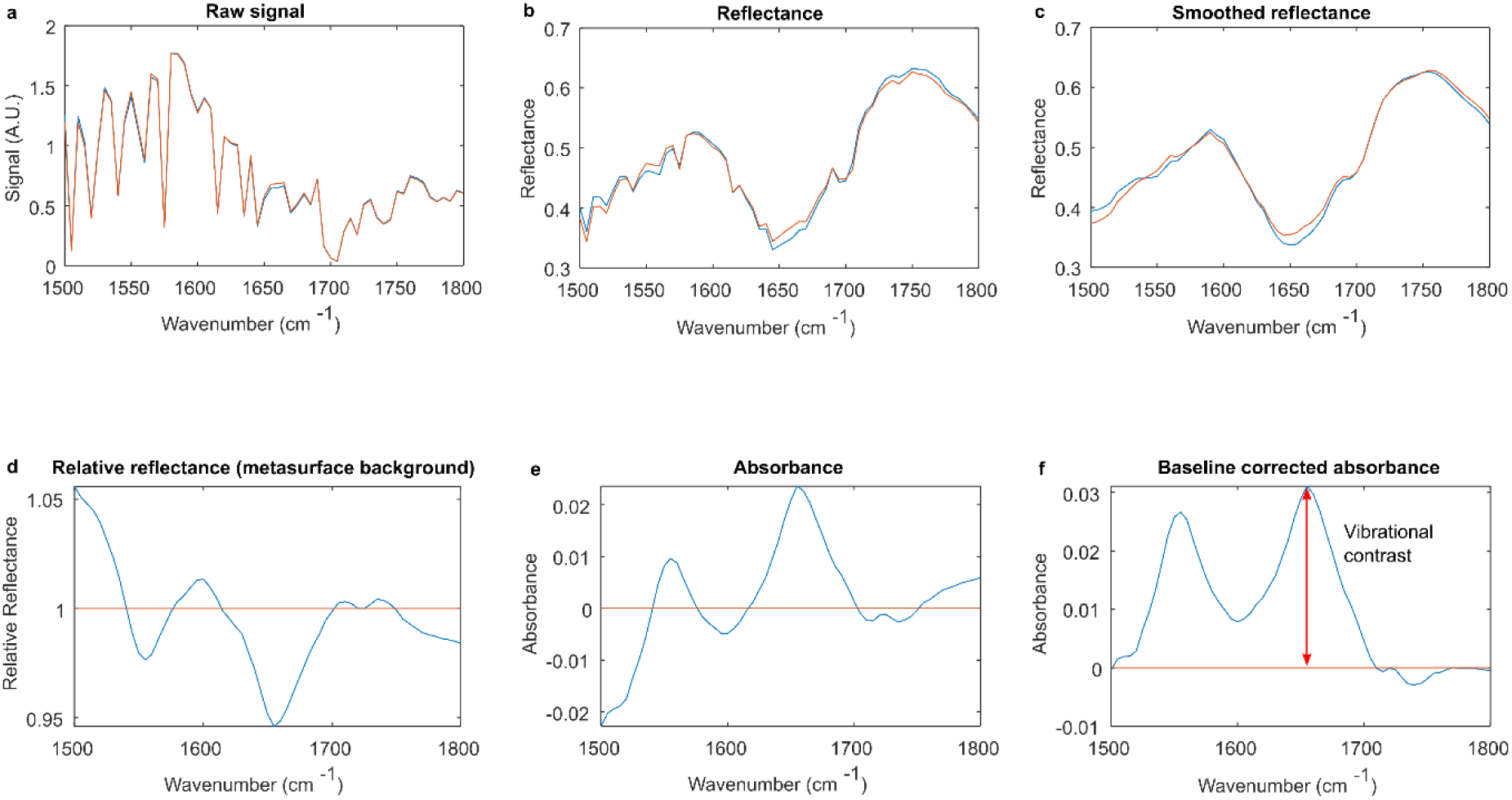
Spectral pre-processing for the protein absorption band (1,500 cm^-1^ – 1,800 cm^-1^). Blue and red curves show the spectrum from cell nucleus and cell-free metasurface background position, as indicated in Figure 3a in the main text. See Methods section in main text for the description of each step. (a) Raw signal from MCT detector. (b) Reflectance spectra at the two positions, using reflectance from a large patch of gold (*i.e.* not nanoantennas) as the background. (c) Reflectance spectra smoothed using Savitzky-Golay filter to remove water vapor related spikes. (d) Relative reflection spectra, using cell-free metasurface as the background. (e) Absorbance spectra computed from the relative reflection spectra. (f) Baseline-corrected absorbance spectra, using a linear baseline correction. The baseline-corrected absorbance was used as the vibrational contrast, as shown in Figure 3 and Figure 4 in the main text.

**Figure S6.**
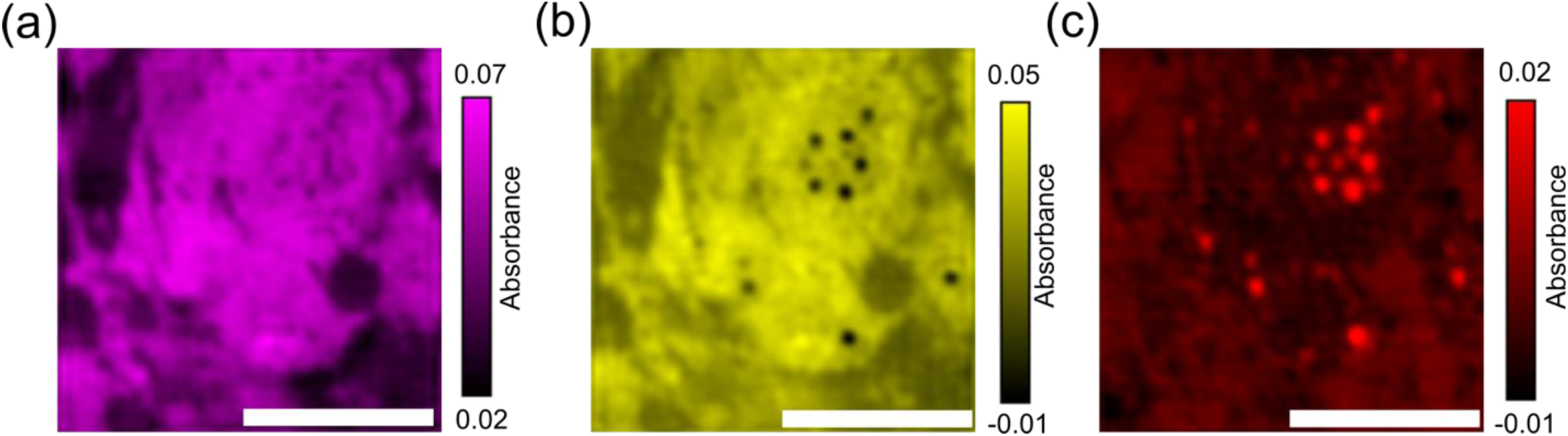
MIRIAM-based MIR image of live adipocyte cells. Discrete frequency MIR images at (a) amide II, (b) amide I, and (c) ester carbonyl C=O bands. Images of live adipocytes (same sample as shown in Figure 4 of main text) were taken just before fixation. Images were taken at 5 discrete frequencies: 1,555 cm^-1^ (amide II), 1,658 cm^-1^ (amide I), 1,740 cm^-1^ (ester carbonyl), as well as 1,502 cm^-1^ and 1,799 cm^-1^ for linear baseline correction. Cell morphology can be seen from the amide II/I vibrations and lipid droplets can be seen from the C=O ester carbonyl vibration. Qualitatively, the image looks very similar to the cell image after fixation (Figure 4 in the main text), though there may have been some cell shrinkage upon fixation. Scale bars: 100 µm.

**Figure S7.**
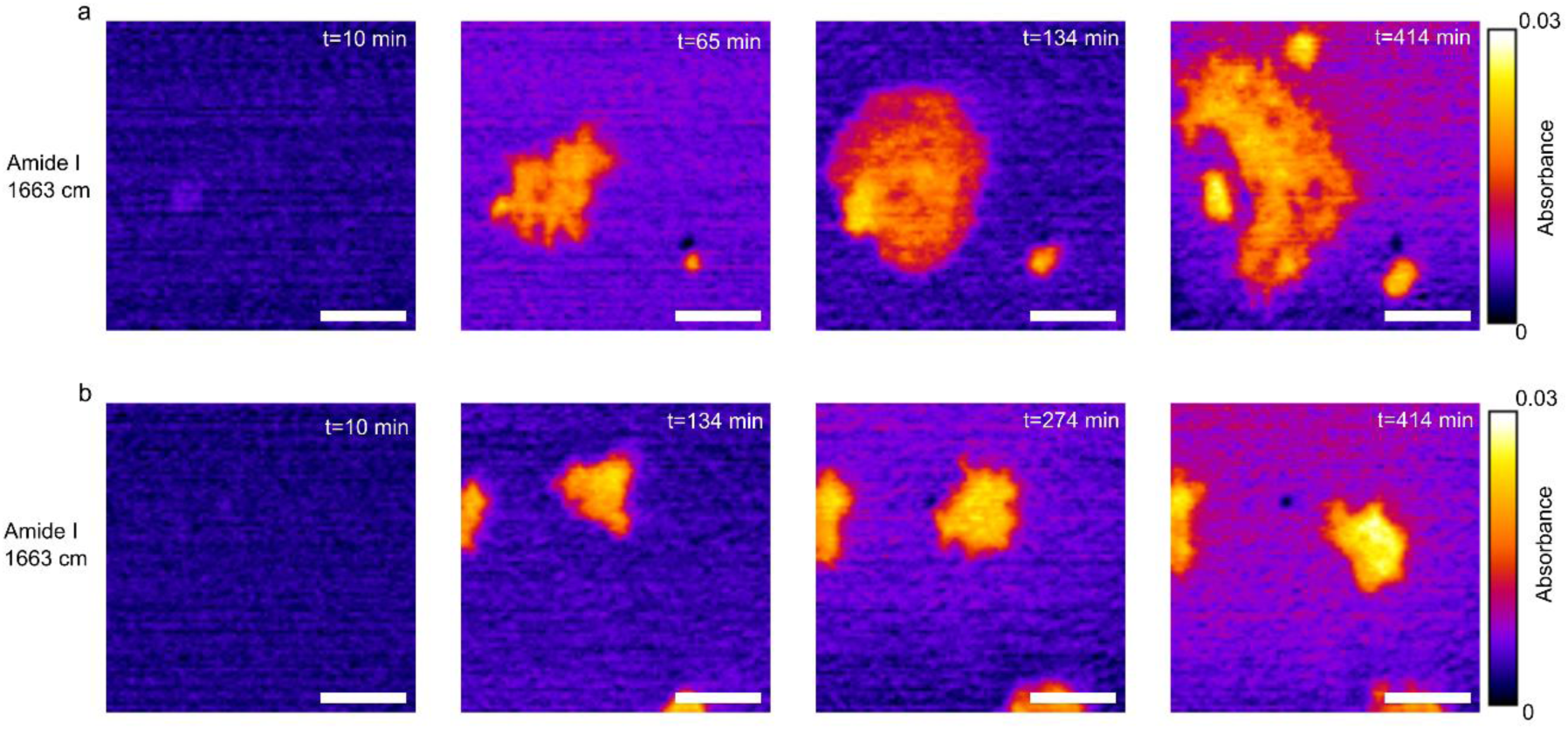
Time-lapse imaging of living 3T3-L1 fibroblasts at 1,663 cm^-1^. Note that because no baseline subtraction was used for these images, contrast here is not purely from amide I vibrational absorption, but there is some contribution from resonance shift mixed in. (a) Adhesion and spreading of a fibroblast cell on the metasurface over 7 hours. (b) Motility of a fibroblast cell on the metasurface. Scale bars: 50 µm.

**Figure S8.**
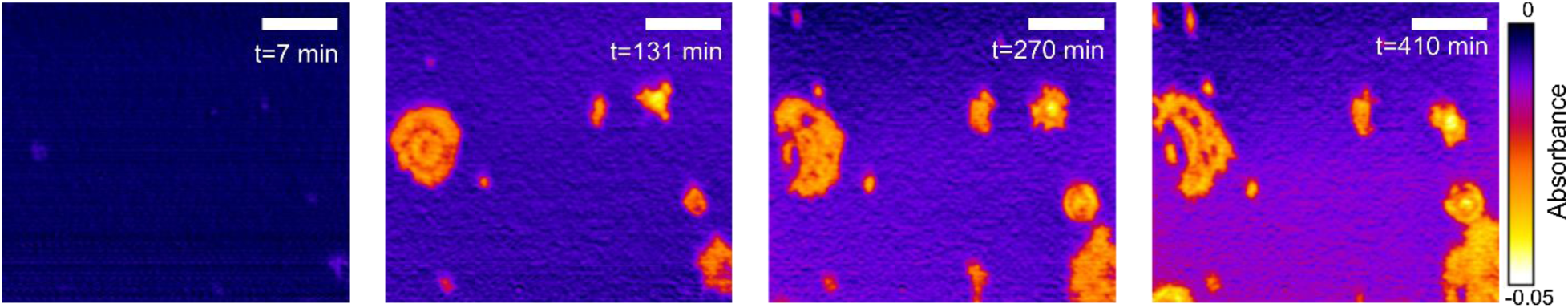
Time-lapse imaging of living 3T3-L1 fibroblasts at 1,502 cm^-1^, full field of view (460 µm × 400 µm). Zoomed in images are shown in Figure 5 of the main text. Scale bars: 100 µm.

**Figure S9.**
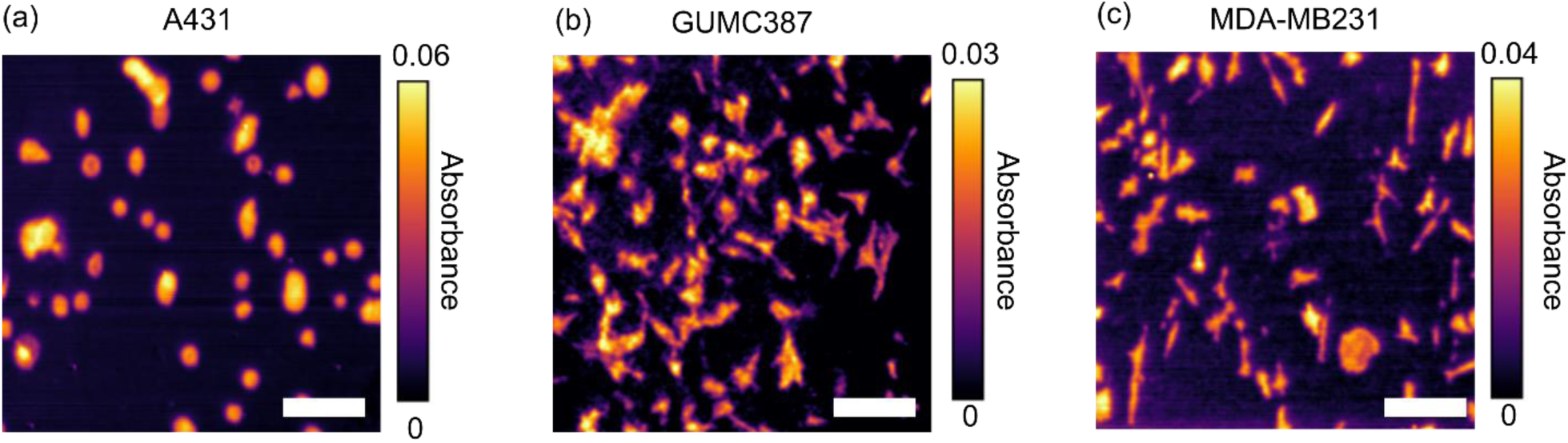
Amide II vibrational contrast images of different cell lines. Images were taken using living cells without fixation. (a) A431 human epidermoid carcinoma cells. (b) GUMC387 conditionally-reprogrammed human prostate cancer cells. (c) MDA-MB231 human breast cancer cells. Different cell morphologies can be seen from each cell line. Scale bar: 100 µm.

## Supplementary Movies

**Movie S1.** Time-lapse imaging of spreading 3T3-L1 fibroblast at 1,502 cm^-1^ (corresponding to Figure 5a). The color mapping is the same as Figure 5a in the main text.

**Movie S2.** Time-lapse imaging of motile 3T3-L1 fibroblast at 1,502 cm^-1^ (corresponding to Figure 5b). The color mapping is the same as Figure 5b in the main text.

**Movie S3.** Time-lapse imaging of spreading 3T3-L1 fibroblast at 1,663 cm^-1^ (corresponding to Figure S7a). The color mapping is the same as Figure S7a in the main text.

**Movie S4.** Time-lapse imaging of motile 3T3-L1 fibroblast at 1,663 cm^-1^ (corresponding to Figure S7b). The color mapping is the same as Figure S7b in the main text.

